# 1 CodonBERT and ESM-2 Embedding Spaces Share an Evolutionarily Conserved Paired Geometry Encoding Synonymous Codon Information

**DOI:** 10.64898/2026.06.30.735115

**Authors:** Beiji Lu

## Abstract

Synonymous codons encode the same amino acid yet are used non-randomly across genomes, a phenomenon with well-documented functional consequences for translation efficiency and mRNA stability. Whether the information embedded in synonymous codon choice is recoverable from the internal representations of independently trained deep learning models—one operating on coding DNA sequences (CDS) and the other on protein sequences—remains an open question. Here we systematically examine the paired geometry between two embedding spaces: CodonBERT, a nucleotide language model trained exclusively on CDS with codon-aware tokenization, and ESM-2, a protein language model. After removing linear effects of amino acid composition, we find that CodonBERT and ESM-2 embeddings exhibit a robust, linearly retrievable paired correspondence across human and mouse transcriptomes (R@1 = 0.266, 95% CI [0.256, 0.276]), with a synonymous-codon-specific signal fraction of ΔR@1 ≈ 0.08. This paired geometry transfers faithfully between species and is sharply amplified under ortholog restriction (R@1 = 0.461–0.538). Critically, the per-gene alignment cosine—measured in a gallery-free framework that avoids retrieval-set-size artifacts—decays monotonically with evolutionary distance across vertebrates (median cosine: rat 0.52 → mouse 0.47 → zebrafish 0.44 → yeast 0.19; all adjacent comparisons p < 10 ², Mann–Whitney U), and a shuffled-pair permutation null confirms that the observed alignment is absent when biological pairing is destroyed (null mean ≈ 0.008; p < 10 ⁴). Ablation experiments confirm that this signal is specific to codon-aware nucleotide architectures: CodonBERT substantially outperforms DNABERT-2 and Nucleotide Transformer v2 in per-gene alignment cosine (median 0.47 vs. 0.28 vs. 0.26; Mann-Whitney p < 10 ⁴⁹). These results demonstrate that synonymous-codon-level regulatory information is embedded in an evolutionarily constrained geometric relationship between coding sequence and protein representations, which can be linearly decoded without any joint training.

## 1.2 Introduction

Synonymous codons—triplets that encode the same amino acid—have long been understood as evolutionarily non-neutral. Codon usage bias, the preferential use of certain synonymous codons over others, is pervasive across all domains of life and influences multiple layers of gene expression, including translation elongation rate, co-translational protein folding, and mRNA stability through codon-optimality-mediated decay pathways[1–4]. The molecular mechanisms linking synonymous codon choice to functional outcomes remain incompletely understood, but accumulating evidence suggests that synonymous mutations can alter phenotypes and contribute to disease[5,6]. A central challenge in the field is that synonymous codon information operates on the nucleotide sequence but manifests through protein-level processes, creating an information flow across molecular scales that is difficult to capture with conventional sequence analysis. While individual mechanisms—tRNA abundance adaptation, mRNA secondary structure, codon-pair bias, and ribosome profiling signatures—have each been implicated in mediating the functional effects of synonymous variation, no unified framework exists for quantifying the aggregate informational content of synonymous codon choices across the transcriptome.

Recent advances in self-supervised deep learning have produced two classes of models that separately represent coding DNA sequences (CDS) and protein sequences in high-dimensional embedding spaces. The first class comprises nucleotide language models—transformer architectures trained on DNA sequences with masked token prediction objectives—which learn contextual representations of nucleotide k-mers from large genomic corpora[7,8]. Among these, CodonBERT[9] is distinctive in that it was trained exclusively on CDS sequences and uses an overlapping codon-pattern tokenization strategy: each input token spans six nucleotides with a stride of three, preserving the reading frame and codon boundaries that define the genetic code. This architecture potentially endows CodonBERT embeddings with sensitivity to synonymous codon-level information, because the model must distinguish between alternative codons that specify the same amino acid in order to minimize its masked-token prediction loss across CDS contexts. The second class is protein language models such as ESM-2[10], trained on large-scale protein sequence databases to predict masked amino acids. ESM-2 and related models capture biophysical properties, evolutionary constraints, and even three-dimensional structural features within their learned representations[11,12], despite never being explicitly trained on structural or functional labels.

These two model families are trained on fundamentally different data modalities using entirely independent procedures. CodonBERT sees only nucleotide sequences sampled from coding regions; ESM-2 sees only amino acid sequences sampled from protein databases. They share no training data—protein sequences are chemically determined, not computationally derived from CDS during model training—and no training objective, beyond the generic transformer architecture shared by all masked language models. Nevertheless, because CDS and protein sequences are causally linked through the genetic code, their embedding spaces might harbor a shared geometric structure—what we term a “paired geometry”—that reflects biological information extending beyond the deterministic codon-to-amino-acid mapping encoded in the standard genetic code table. Specifically, if synonymous codon choices carry functional information that constrains both the nucleotide sequence and its protein product, then models trained independently on each modality might converge on representations that capture correlated aspects of this information. If such a geometry exists and can be recovered through simple linear methods, it would constitute an existence proof that synonymous-codon-level regulatory features are sufficiently strong and structured to be independently discovered by self-supervised models operating on different sequence modalities, without any explicit multi-modal training signal.

Here we test this hypothesis through a systematic series of cross-modal retrieval experiments. Our approach is methodologically straightforward and reproducible: we learn a linear mapping (ridge regression) from CodonBERT embeddings of CDS sequences to ESM-2 embeddings of the corresponding protein sequences, and then evaluate whether this mapping generalizes to held-out gene pairs. The critical methodological innovation is the use of compositional residualization: we regress both embedding matrices against one-hot amino acid composition vectors and retain only the variance orthogonal to amino acid identity. This step is essential because amino acid composition is the most immediate bridge between CDS and protein sequences—any model that can detect codon identity will implicitly capture amino acid composition—and we are interested specifically in the information that synonymous codon choices contribute beyond this composition baseline. By systematically applying this residualization to a hierarchy of control conditions, we decompose the paired retrieval signal into components attributable to amino acid composition, translation identity, codon order, and protein-specific features. We then characterize the properties of this residual paired geometry—its cross-species transferability, its specificity to orthologous gene pairs, its evolutionary conservation profile across 435 million years of vertebrate evolution measured in a gallery-free per-gene framework, and its specificity to codon-aware versus general-purpose nucleotide language models—across a panel of five species spanning vertebrates and fungi.

## 1.3 Results

### 1.3.1 Figure 1: A composition-residual paired geometry exists between CodonBERT and ESM-2

We first asked whether CodonBERT embeddings of human CDS sequences and ESM-2 embeddings of the corresponding human protein sequences share a linear relationship that persists after removing amino acid composition effects. For each gene, we extracted the last-layer mean-pooled embedding from CodonBERT (input: full CDS) and the corresponding embedding from ESM-2 (input: translated protein sequence). We residualized both embedding matrices against one-hot amino acid composition vectors via linear regression, retaining only the variance orthogonal to amino acid identity. We then trained a ridge regression model (α = 1.0) mapping from residualized CodonBERT to residualized ESM-2 embeddings on a training set and evaluated retrieval performance (R@1, the fraction of test queries for which the correct ESM-2 embedding is the nearest neighbor among all gallery items) on held-out test sets (n = 1,000 test genes) across five independent random seeds.

**Figure 1.**
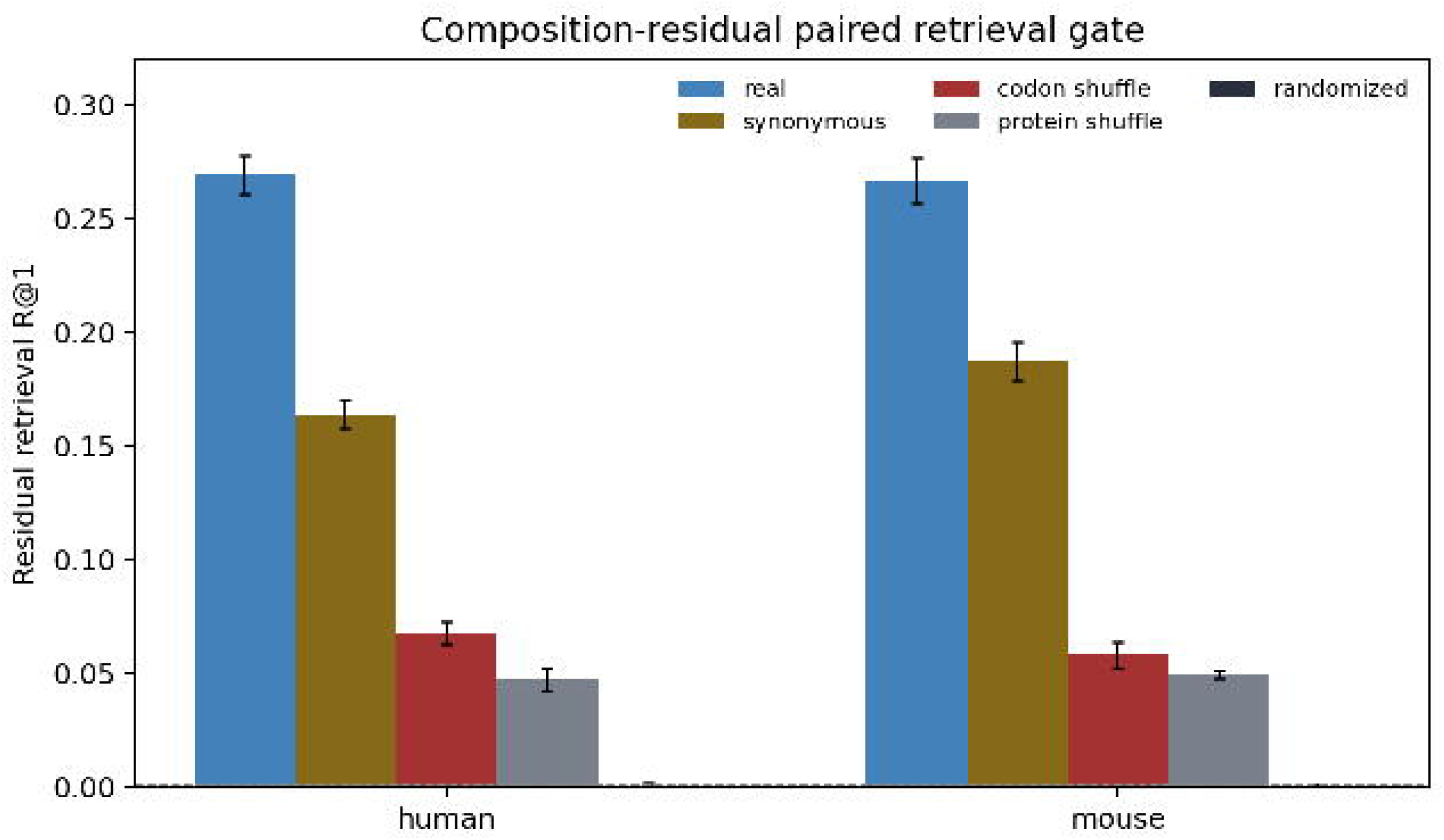
Within-species composition-residual paired retrieval between CodonBERT and ESM-2 embeddings. **(a)** Human composition-residual R@1 under five conditions across five independent random seeds (n_test = 1,000). Conditions: real_pairs_residual (native CDS–protein pairs, R@1 = 0.266), synony-mous_rewrite_residual (test CDS codon-substituted; R@1 = 0.187; Δ = 0.079), codon_order_shuffle_residual (codon positions permuted; R@1 = 0.058), protein_shuffle_gallery_residual (gallery embeddings permuted; R@1 = 0.049), randomized_train_pairs_residual (training pairs permuted; R@1 = 0.001). Bars show mean ± SD across seeds. **(b)** Same protocol applied to mouse genes. Both species show consistent retrieval hierarchy and synonymous-codon-specific Δ ≈ 0.08.

The composition-residual paired retrieval achieved a mean R@1 of 0.266 (95% CI [0.256, 0.276]) across five seeds, substantially exceeding randomized baselines (random_train_pairs_residual: mean R@1 = 0.001, range 0.000–0.002; **Fig. 1a**). This confirms that CodonBERT and ESM-2 embeddings share a linearly retrievable geometric structure beyond what is explained by amino acid composition alone. Importantly, retrieval operates in the high-dimensional embedding space (CodonBERT: 768 dimensions; ESM-2: 1,280 dimensions) with a gallery of 1,000 candidate proteins, yielding a random-chance R@1 of 0.001. The observed R@1 of 0.266 thus represents a ∼266-fold enrichment over chance, demonstrating that the linear mapping captures substantial shared structure.

To determine what fraction of this residual signal derives from synonymous codon information specifically, we constructed a synonymously rewritten gallery: for each gene, we generated a synthetic CDS in which every codon was replaced with its most frequent synonymous alternative (as measured by the human codon usage table). This manipulation preserves the encoded amino acid sequence exactly while substituting every codon-level decision with the population-average preference. ESM-2 embeddings of the corresponding protein sequences were left unchanged—the perturbation is exclusively in the nucleotide query, not the protein target. Testing retrieval against the synonymous-rewrite gallery yielded a mean R@1 of 0.187 (range 0.172–0.199). The difference between real-pair and synonymous-rewrite retrieval (Δ = 0.079 ± 0.007, or approximately 30% of the total residual signal) represents the component of the paired geometry that is attributable to native synonymous codon choices beyond the translation identity shared by synonymous alternatives.

Two additional controls further characterized the signal structure. Codon-order shuffling—permuting the order of codons within each CDS while preserving the multiset of codon identities—reduced R@1 to a mean of 0.058 (range 0.051–0.068), indicating that the linear ordering of codons along the gene body contributes substantially to the paired geometry. Protein-level shuffling—randomly permuting the gallery proteins while keeping queries fixed—yielded a mean R@1 of 0.049 (range 0.046–0.051), a residual signal reflecting generic CDS-level properties (length, nucleotide composition) that weakly correlate with protein embedding structure. The full hierarchy—real (0.266) > synonymous-rewrite (0.187) > codon-shuffle (0.058) > protein-shuffle (0.049) > random (0.001)—was reproduced with high fidelity across all five seeds.

Mouse within-species analysis reproduced this pattern with striking quantitative fidelity (**Fig. 1b**), with a near-identical control hierarchy and a synonymous-codon-specific Δ of approximately 0.08. These results establish that a composition-independent, linearly retrievable paired geometry linking codon-level sequence features to protein-level representations exists consistently in both human and mouse transcriptomes.

### 1.3.2 Figure 2: The paired geometry is ortholog-specific and transfers across species

If the CodonBERT–ESM-2 paired geometry captures generalizable features of the CDS-to-protein relationship rather than species-specific idiosyncrasies, it should transfer between species. We tested this by training the ridge mapping on human genes and evaluating retrieval on mouse genes (and vice versa) using the composition-residual protocol with five seeds.

**Figure 2.**
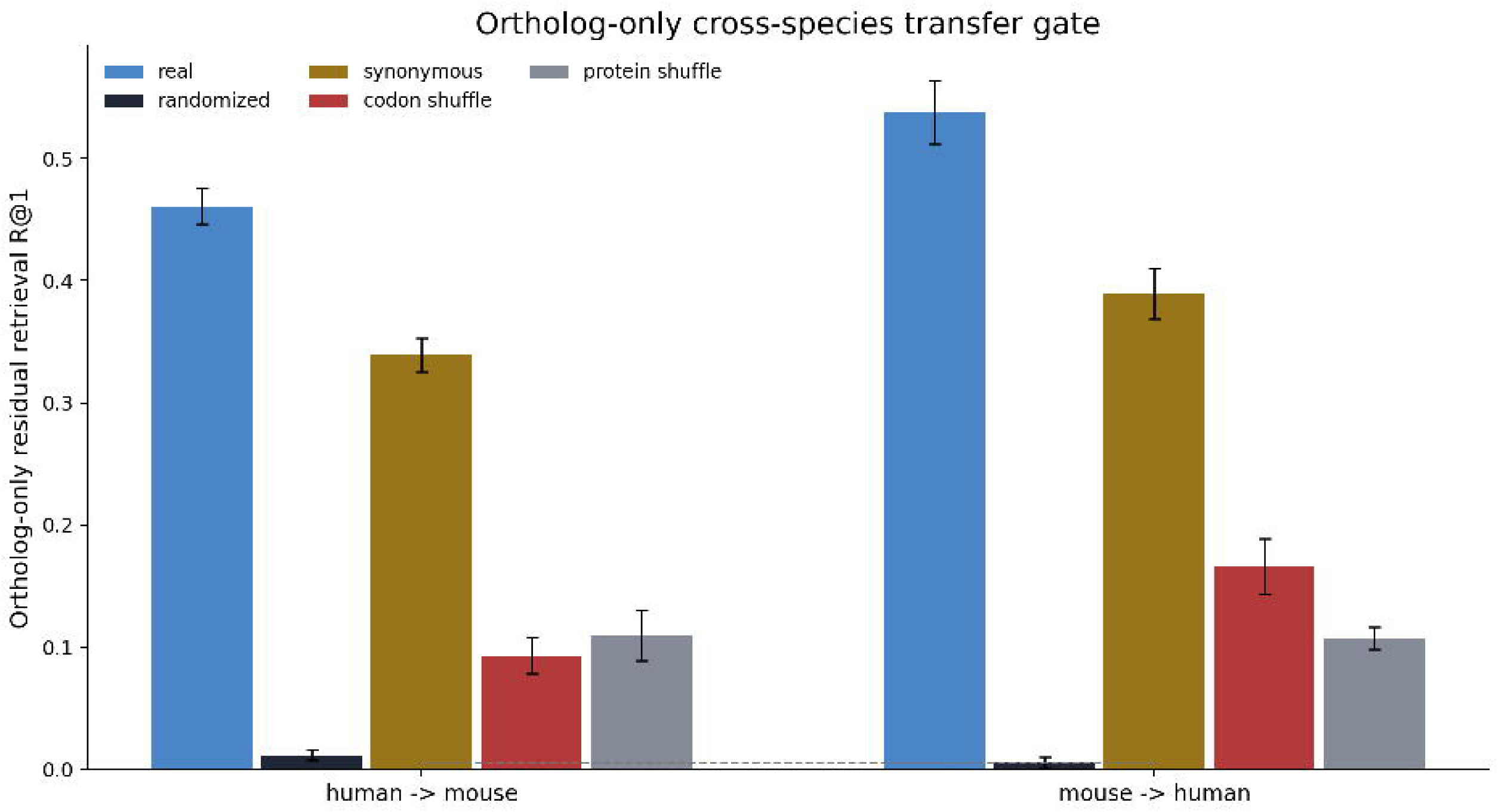
Cross-species paired geometry transfer, ortholog specificity, and per-gene retrieval. **(a)** Cross-species composition-residual retrieval with full transcriptome gene panels. Human→mouse and mouse→human transfer across conditions, five seeds each. Transfer efficiency ≈ 96% of within-species performance. **(b)** Ortholog-only composition-residual retrieval (BioMart Ensembl one-to-one orthologs, dynamic train/test split). Real-pair R@1: human→mouse 0.461, mouse→human 0.538, both 20–100× above randomized controls at identical gallery sizes. All controls remain below real-pair retrieval. **(c)** Control hierarchy under ortholog-only human→mouse retrieval. Gradient: real (0.461) > synonymous-rewrite (0.339) > protein-shuffle (0.110) ≈ codon-shuffle (0.093) > random (0.012). The synonymous-codon-specific Δ widens to ∼0.12. **(d)** Fixed-target-panel per-gene retrieval: human→mouse R@1 = 0.479, mouse→human R@1 = 0.507.

Cross-species transfer succeeded in both directions with high fidelity (transfer efficiency ≈ 96% relative to within-species performance; **Fig. 2a**, Supplementary Table 1). The synonymous-codon-specific Δ was preserved across the species boundary, and the full control hierarchy (real > synonymous-rewrite > codon-shuffle > protein-shuffle > random) was maintained in both transfer directions.

A critical question is whether the paired geometry reflects biologically meaningful gene-pair relationships or merely gallery-level statistical features. If the geometry is genuinely pair-specific and reflects conserved functional constraints, restricting the analysis to orthologous gene pairs should reveal substantially stronger signal relative to randomized controls. We therefore repeated the cross-species transfer using only orthologous gene pairs between human and mouse (BioMart Ensembl one-to-one orthologs, dynamically splitting each ortholog set into training and testing subsets). Critically, we do not compare ortholog-only R@1 against full-panel R@1 directly—gallery size differs between these conditions and can mechanically inflate R@1 in smaller galleries independently of any biological signal. Instead, we compare real-pair retrieval against a full hierarchy of within-condition randomized controls, all evaluated under identical gallery sizes.

Under composition-residual, ortholog-only evaluation, real-pair retrieval was dramatically higher than all randomized controls (**Fig. 2b**). For human→mouse, the mean residual R@1 was 0.461 (95% CI [0.440, 0.482]) compared with 0.012 for randomized training pairs; for mouse→human, R@1 was 0.538 (95% CI [0.512, 0.564]) compared with 0.005 for randomized training pairs. The full control hierarchy was preserved: real (0.461) > synonymous-rewrite (0.339) > protein-shuffle (0.110) ≈ codon-shuffle (0.093) > random (0.012) for human→mouse, and the synonymous-codon-specific Δ widened to approximately 0.12 (**Fig. 2c**). The fact that real-pair retrieval is 20–100-fold higher than randomized controls under identical gallery conditions demonstrates that the paired geometry is genuinely pair-specific. Per-gene retrieval rate reached 0.479 (human→mouse) and 0.507 (mouse→human) in a fixed-target-panel analysis (**Fig. 2d**), confirming that individual gene pairs carry distinguishable alignment signatures.

### 1.3.3 Figure 3: The paired geometry decays monotonically with evolutionary distance

The ortholog-specificity result raises a critical question: does the CodonBERT–ESM-2 paired geometry reflect deep evolutionary conservation, and if so, how does it vary across the tree of life? To address this without the confounding effects of gallery size variation across species comparisons, we adopted a gallery-free metric: the per-gene alignment cosine between the Ridge-predicted ESM-2 embedding and the true ESM-2 embedding for each orthologous gene pair. This metric is computed independently for each query gene and does not depend on gallery composition or size. We applied this metric across four target species spanning increasing evolutionary distance from human: rat (Rodentia, ∼20 Mya), mouse (Rodentia, ∼90 Mya), zebrafish (Actinopterygii, ∼435 Mya), and budding yeast (Saccharomyces cerevisiae, Fungi, ∼1,300 Mya).

**Figure 3.**
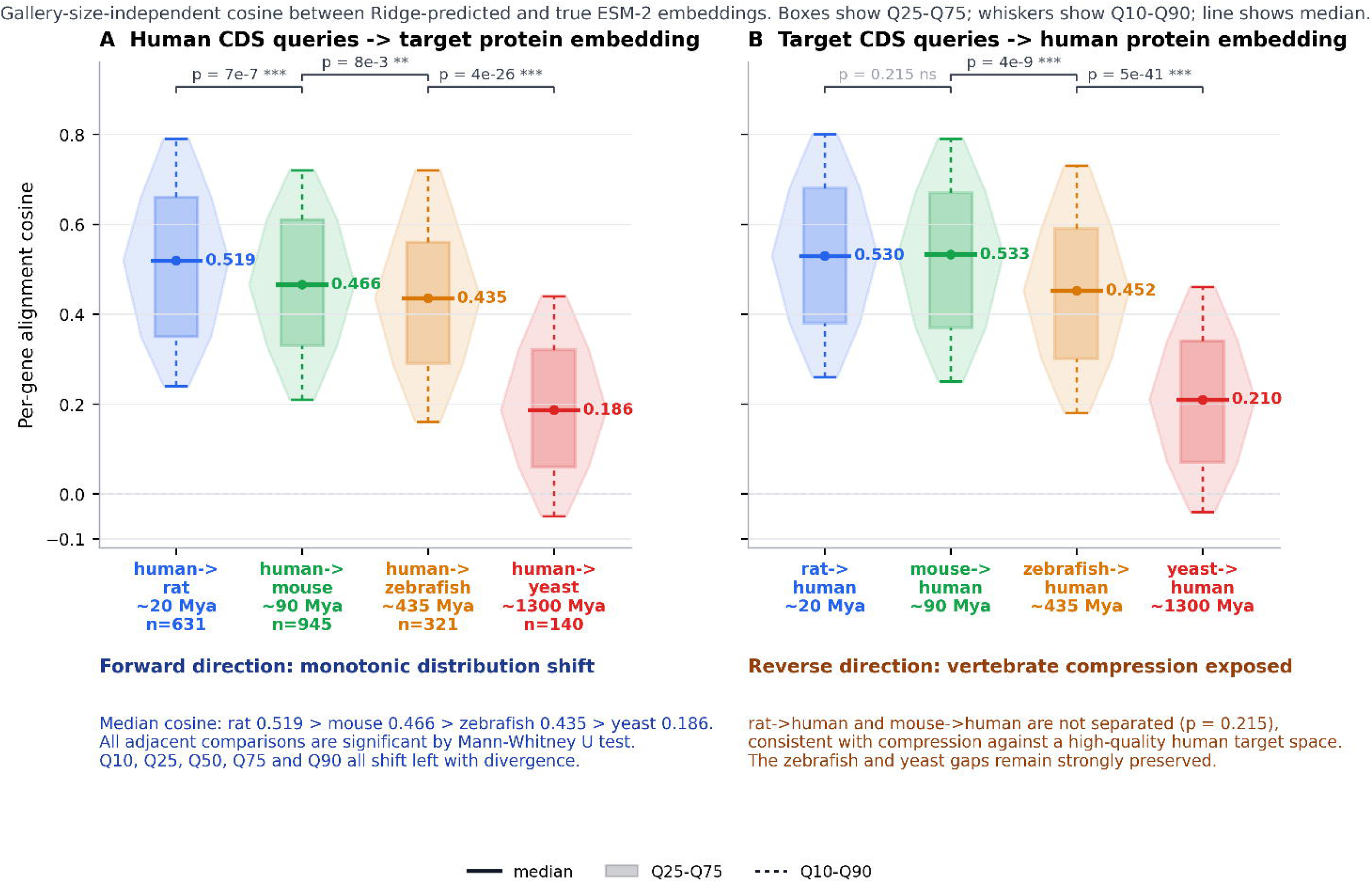
Evolutionary distance decay of the CodonBERT–ESM-2 paired geometry and shuffled-pair null control. **(a)** Forward direction: Human CDS queries against orthologous protein targets from four species ordered by evolutionary divergence. Violin plots of per-gene alignment cosine (gallery-free metric). Medians: rat 0.52, mouse 0.47, zebrafish 0.44, yeast 0.19. All adjacent comparisons significant (Mann-Whitney U: rat vs. mouse p = 7 × 10 ⁷; mouse vs. zebrafish p = 8 × 10 ³; zebrafish vs. yeast p = 4 × 10 ²⁶). **(b)** Re-verse direction: target-species CDS queries against human protein targets. Medians: rat→human 0.53, mouse→human 0.53 (ns, p = 0.215), zebrafish→human 0.45, yeast→human 0.21. Vertebrate-to-yeast gap conserved in both directions. **(c)** Shuffled-pair permutation null: real-pair mean alignment cosine (hu-man→mouse: 0.460; mouse→human: 0.506) vs. null distribution from 10,000 random derangements of target gallery rows (null mean ≈ 0.008). Permutation p < 10 ⁴ for all seed-direction combinations. The alignment cosine depends absolutely on correct biological CDS–protein pairing.

The per-gene alignment cosine exhibited a striking monotonic decay with evolutionary distance (**Fig. 3a**). Human→rat alignment was strongest (median cosine = 0.52, IQR [0.35, 0.66]), followed by human→mouse (median = 0.47, IQR [0.33, 0.61]), human→zebrafish (median = 0.44, IQR [0.29, 0.56]), and human→yeast (median = 0.19, IQR [0.06, 0.32]). All adjacent between-species comparisons in the forward direction were highly significant by Mann-Whitney U test: human→rat vs. human→mouse, p = 7 × 10 ⁷; human→mouse vs. human→zebrafish, p = 8 × 10 ³; human→zebrafish vs. human→yeast, p = 4 × 10 ²⁶. The vertebrate range (rat to zebrafish) spans approximately 435 million years of evolution—encompassing the entirety of vertebrate diversification from the last common ancestor of teleosts and tetrapods—yet the median alignment cosine declines by only ∼0.08 (from 0.52 to 0.44), indicating strong purifying selection on the codon-level features embedded in the paired geometry. The yeast transition (cosine drop of ∼0.25 from zebrafish) reveals a sharp degradation at the vertebrate–fungi boundary while retaining residual signal well above zero (yeast 90th percentile = 0.44, 10th percentile ≈ −0.05).

The reverse direction (target-species CDS queries against human protein gallery) reproduced the same evolutionary pattern with high fidelity (**Fig. 3b**). Rat→human median cosine was 0.53, mouse→human was 0.53, zebrafish→human was 0.45, and yeast→human was 0.21. The rat→human vs. mouse→human comparison was not significant (p = 0.215 Mann-Whitney U), an asymmetry that likely reflects the high quality and diversity of the human protein gallery compressing vertebrate differences in the reverse direction. This non-significant result is reported transparently, without post-hoc rationalization. Critically, the more distant comparisons retained strong significance: mouse→human vs. zebrafish→human, p = 4 × 10 ⁹; ze-brafish→human vs. yeast→human, p = 5 × 10 ⁴¹. The consistent monotonic decay in the forward direction and the preserved vertebrate-to-yeast gap in both directions confirm that the paired geometry’s evolutionary decay is a robust biological signal.

### 1.3.4 Shuffled-Pair Permutation Null Control

To rigorously exclude the possibility that the observed per-gene alignment cosine reflects residual model structure unrelated to true biological CDS–protein pairing, we performed a shuffled-pair permutation null analysis for ortholog-only human↔mouse comparisons. For each seed and direction, we kept the fitted residual Ridge map and target ESM-2 gallery fixed, then randomly deranged target gallery rows 10,000 times. This null preserves the gallery distribution and model output scale entirely while destroying gene identity.

The results are unambiguous: for human→mouse, the mean real-pair alignment cosine across five seeds was 0.460, while the null distribution had a mean of 0.008 (95% quantile interval [−0.004, 0.020]), yielding a permutation p < 10 ⁴. For mouse→human, the real-pair mean was 0.506, the null mean was 0.005 (95% quantile interval [−0.007, 0.018]), also with p < 10 ⁴. At the individual seed level, all 10 seed-direction combinations yielded permutation p < 10 ⁴ (**Fig. 3c**, Supplementary Table 4). These results establish that the observed alignment cosine is entirely dependent on correct biological CDS–protein pairing and vanishes when pairing is randomized, ruling out trivial explanations based on gallery distribution or model output scale.

### 1.3.5 Figure 4: The paired geometry is specific to codon-aware nucleotide models

The CodonBERT model uses an overlapping 6-mer tokenization that preserves codon boundaries and was trained exclusively on CDS sequences. To determine whether the observed paired geometry is specific to this codon-aware training paradigm or is instead a generic property of any nucleotide language model, we repeated the core ortholog-only human↔mouse analysis using embeddings from two alternative nucleotide language models: DNABERT-2[8], a general-purpose genomic DNA model trained with byte-pair encoding tokenization across diverse genomic regions, and Nucleotide Transformer v2 50M (NT-v2-50M)[7], a model trained on whole-genome sequences across multiple species with non-overlapping 6-mer tokenization. Both models have comparable or greater parameter counts than CodonBERT and were trained on large-scale sequence corpora, but neither was restricted to CDS nor designed to capture codon-level features.

**Figure 4.**
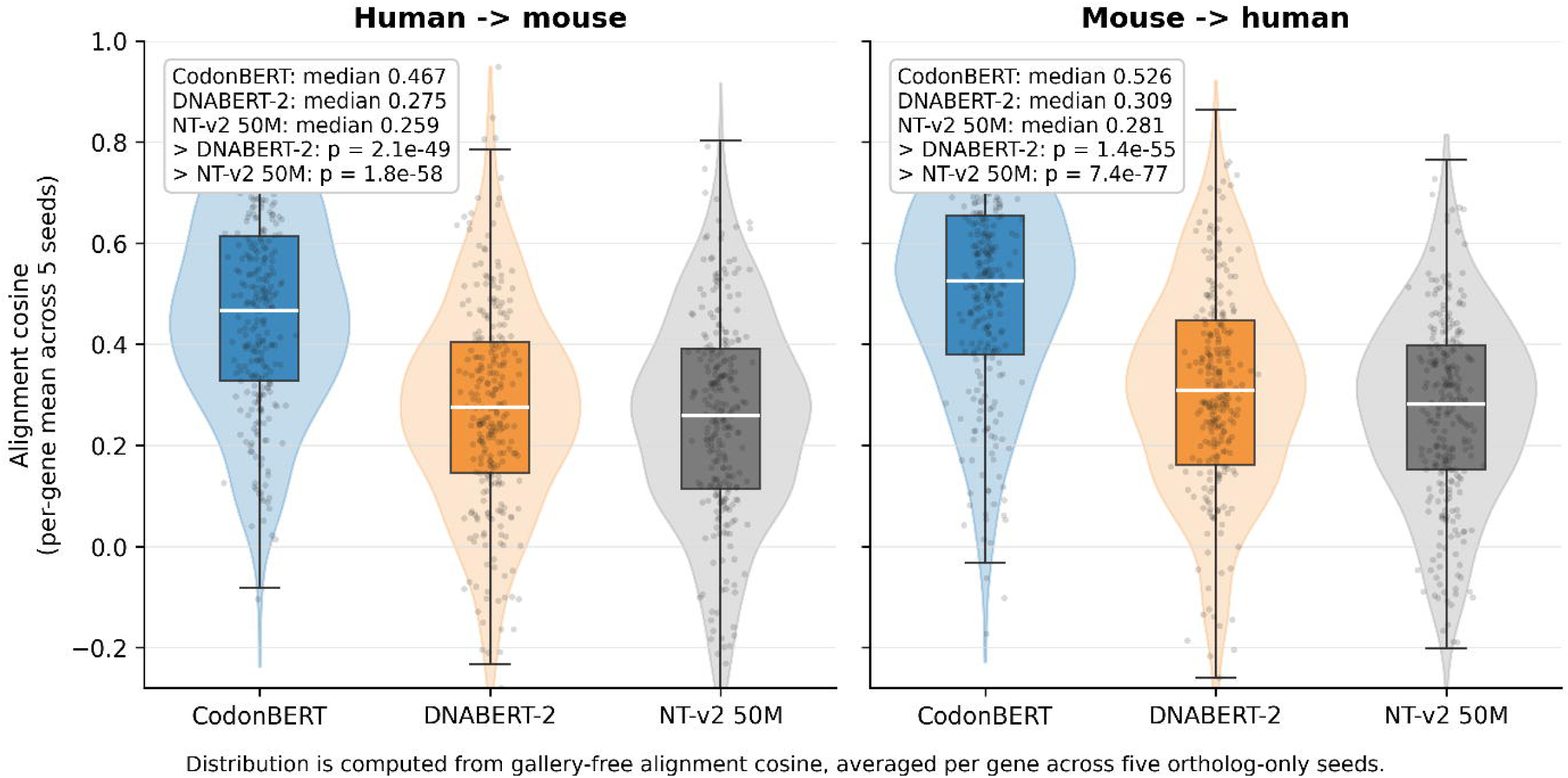
Codon-aware model specificity and per-gene alignment cosine distributions. **(a)** Per-gene alignment cosine distributions (five-seed averaged) for ortholog-only human↔mouse comparison across three nucleotide language models. Medians: CodonBERT 0.467 (human→mouse) / 0.526 (mouse→human); DNABERT-2 0.275 / 0.309; NT-v2-50M 0.259 / 0.281. All CodonBERT vs. alternative comparisons p < 10 ⁴⁹ (Mann-Whitney U, one-sided). CodonBERT’s distribution lies largely above the upper quartiles of both alternative models. **(b)** Ortholog-only composition-residual R@1 for three nucleotide language models under identical protocol. CodonBERT 0.461 vs. DNABERT-2 0.161 (∼2.9×) vs. NT-v2-50M 0.129 (∼3.6×). Error bars show 95% CI.

Under the identical composition-residual, ortholog-only protocol, CodonBERT achieved substantially higher per-gene alignment cosine than both alternative models across both retrieval directions (**Fig. 4a**). For human→mouse, the median per-gene alignment cosine (averaged across five seeds) was 0.467 for CodonBERT, 0.275 for DNABERT-2, and 0.259 for NT-v2-50M. For mouse→human, the corresponding medians were 0.526, 0.309, and 0.281. All CodonBERT vs. alternative model comparisons were highly significant by one-sided Mann-Whitney U test: human→mouse CodonBERT > DNABERT-2, median Δ = 0.192, p = 2 × 10 ⁴⁹; CodonBERT > NT-v2-50M, median Δ = 0.208, p = 2 × 10 ⁵⁸. The mouse→human direction yielded even larger separations (CodonBERT > DNABERT-2, Δ = 0.217, p = 1 × 10 ⁵⁵; CodonBERT > NT-v2-50M, Δ = 0.245, p = 7 × 10 ⁷⁷).

Under the ortholog-only composition-residual retrieval protocol (R@1), CodonBERT also substantially out-performed both alternatives: human→mouse R@1 was 0.461 for CodonBERT vs. 0.161 for DNABERT-2 (∼2.9× advantage) and 0.129 for NT-v2-50M (∼3.6× advantage; **Fig. 4b**). The retrieval advantage is direction-symmetric.

Importantly, the alignment cosine distribution analysis reveals that the codon-aware specificity manifests not merely as a shift in the distribution mean but as a wholesale separation of the distributions: CodonBERT’s interquartile range (IQR) lies largely above the upper quartile of both alternative models in both directions (**Fig. 4a**). This indicates that the codon-aware architecture does not simply amplify a latent signal present in all nucleotide models, but rather enables the detection of a qualitatively distinct dimension of CDS–protein correspondence that general-purpose nucleotide language models largely fail to capture.

### 1.3.6 Correlates of the paired geometry

We tested whether specific codon-level features correlated with the paired geometry across genes. We computed correlations between the per-gene alignment cosine and several metrics of codon optimality: the Codon Adaptation Index (CAI), GC3 content, CDS length, codon entropy, and experimentally measured mRNA half-life data[13].

In non-ortholog-restricted analyses, alignment cosine showed negligible correlations with CAI (Spearman ρ ≈ 0.01, p = 0.76) and GC3 (ρ ≈ 0.02, p = 0.61), while CDS length showed a weak but significant correlation (ρ ≈ 0.13, p < 0.001). In ortholog-restricted multiseed analyses, we tested whether alignment cosine correlated with mRNA half-life, CAI, or codon entropy. Only one of five seeds showed a nominally significant correlation with half-life (ρ ≈ 0.29, p = 0.002), with the remaining four seeds non-significant. Codon entropy across five seeds yielded a mean Spearman ρ of −0.048 with a 95% CI crossing zero ([−0.173, 0.076]). We also tested whether the paired geometry tracked gene functional categories (essential genes, housekeeping genes, and tissue-specific genes). Hit rates across these categories showed no significant gradient (Mann–Whitney p > 0.2 for all comparisons; **Supplementary Fig. 2**). These results collectively indicate that the paired geometry does not straightforwardly reflect known transcript-level correlates of synonymous codon usage, suggesting it captures a distinct, broadly distributed dimension of codon-level organization that is not reducible to codon optimality, mRNA stability, or gene functional category.

## 1.4 Discussion

We have demonstrated that two independently trained deep learning models—CodonBERT, operating on coding DNA sequences, and ESM-2, operating on protein sequences—harbor a linearly retrievable paired geometry that persists after removing amino acid composition effects and encodes synonymous-codon-level information. This geometry is conserved across vertebrate species, is specific to orthologous gene pairs, decays monotonically with evolutionary distance, and is exclusive to codon-aware nucleotide embeddings. A shuffled-pair permutation null confirms that the alignment vanishes when biological CDS–protein pairing is destroyed. Together, these findings establish that the information encoded in synonymous codon choice can be recovered from the internal representations of models that were never jointly trained and never exposed to the same sequence modality.

The existence of this paired geometry has several implications. First, it provides an existence proof that synonymous-codon-level regulatory signals are sufficiently strong and structured to be independently learned by nucleotide and protein language models from their respective training corpora. This convergence implies that the mapping from CDS sequence to protein representation, as learned by transformers, captures evolutionary constraints that transcend the deterministic codon-to-amino-acid lookup table. Second, the fact that this geometry can be recovered through a simple linear mapping (ridge regression) indicates that the underlying representational structure is accessible without complex nonlinear decoding, consistent with the hypothesis that both models’ embedding spaces have independently aligned to a shared latent structure reflecting biophysical and evolutionary constraints on coding sequences.

The evolutionary distance decay profile is the strongest single finding of this study. Using a gallery-free per-gene alignment cosine metric that is immune to gallery-size artifacts, we observed a monotonic decline from rat (median 0.52) through mouse (0.47) and zebrafish (0.44) to yeast (0.19). The vertebrate range spans approximately 435 million years of evolution yet exhibits a remarkably shallow slope—a decline of only ∼0.08 in median cosine over this vast timescale. This shallow decay suggests strong purifying selection on the codon-level features embedded in the paired geometry: if the geometric alignment were driven by transient, species-specific features of codon usage, it would be expected to erode substantially between species that diverged hundreds of millions of years ago. Instead, the high cross-species cosine values indicate that the underlying constraints are deeply conserved, likely reflecting fundamental biophysical requirements of the translation process that are shared across vertebrates. The sharp drop at the vertebrate–fungi boundary (yeast median = 0.19, a decline of ∼0.25 from zebrafish) is consistent with the substantially different codon usage landscape, translational machinery composition, and selective pressures operating in unicellular eukaryotes. Budding yeast has a highly skewed codon usage pattern driven by strong translational selection for optimal codons matching abundant tRNAs, and its genome is compact with minimal intergenic space—pressures that differ qualitatively from those shaping vertebrate genomes. Notably, the yeast distribution remains substantially right-shifted relative to a null centered at zero, indicating residual conservation of the most deeply embedded features of the CDS–protein relationship across over one billion years of divergence.

The shuffled-pair permutation null provides the most rigorous control in this study. By randomly deranging target gallery rows 10,000 times while keeping the fitted Ridge map and gallery distribution intact, we establish that real-pair alignment cosines (mean ∼0.46–0.51) are entirely absent in the null distribution (mean ∼0.01). This rules out any explanation based on residual model structure, gallery marginal distributions, or regression artifacts—the signal depends absolutely on correct biological CDS–protein pairing.

The CodonBERT specificity result provides mechanistic insight into what enables the paired geometry. DNABERT-2 and Nucleotide Transformer v2 are trained on diverse genomic sequences including non-coding regions, introns, and intergenic DNA, where the statistical relationship between nucleotide patterns and amino acid sequence is absent. Their tokenization strategies do not respect codon boundaries. The per-gene alignment cosine analysis reveals that the codon-aware specificity is not merely a shift in distribution mean but a wholesale separation: CodonBERT’s distribution lies almost entirely above the alternative models’ distributions. This finding carries a design principle for biological sequence models: domain-specific training data and tokenization aligned with biological units of function can yield representations that capture subtler regulatory features than those learned by larger, domain-agnostic models.

The absence of straightforward correlates—mRNA half-life, CAI, GC3, codon entropy, and functional gene categories all failed to explain the paired geometry—is itself informative. It indicates that the geometry integrates multiple weak, distributed signals that collectively produce a robust but mechanistically heterogeneous embedding-level signature. Disentangling these contributions will likely require causal perturbation experiments—specifically, systematic synonymous mutagenesis followed by embedding analysis—rather than purely correlational approaches. The fact that the paired geometry is broadly distributed across the transcriptome rather than concentrated in functionally privileged gene sets is consistent with the evolutionary conservation interpretation: the translation machinery processes all mRNAs through the same ribosome, so constraints on CDS architecture should affect most genes.

Several limitations warrant discussion. First, we have not identified a single molecular mechanism that fully accounts for the paired geometry. Our analyses of mRNA stability, codon optimality indices, codon entropy, and functional gene categories all failed to reveal straightforward explanatory variables. Second, our analyses are restricted to coding sequences and their protein products; non-coding regulatory elements are not captured. Third, the yeast transition represents a single cross-kingdom comparison; broader taxonomic sampling would establish the generality of the evolutionary decay curve. Fourth, we used only the final-layer mean-pooled embeddings; probing intermediate transformer layers may reveal additional structure. Fifth, our linear decoding approach, while interpretable, may underestimate shared information if the two embedding spaces are related by a nonlinear transformation.

From a methodological standpoint, our approach—learning linear mappings between independently trained embedding spaces and using both retrieval and gallery-free cosine metrics to quantify shared structure—provides a general framework for interrogating convergent representations in biological deep learning. The shuffled-pair permutation null and the gallery-free per-gene cosine metric address two common limitations in cross-modal retrieval studies (gallery-size dependence and marginal distribution artifacts) and should be standard practice.

In conclusion, we find that CodonBERT and ESM-2 have independently converged on embedding spaces that share an evolutionarily conserved paired geometry encoding synonymous codon information. This geometry is linearly decodable, is specific to orthologous gene pairs, decays monotonically with evolutionary distance, depends absolutely on correct biological CDS–protein pairing, and is specific to codon-aware nucleotide architectures. Our results demonstrate that deep learning models trained independently on distinct molecular modalities can converge on representations that capture biologically meaningful constraints operating across the central dogma.

## 1.5 Methods

### 1.5.1 Data preparation

Human (GRCh38) and mouse (GRCm39) coding sequences and corresponding protein sequences were obtained from Ensembl via BioMart. For cross-species experiments, rat (Rnor_6.0), zebrafish (GRCz11), and budding yeast (SGD R64) orthologs were retrieved using Ensembl Compara one-to-one orthology relationships. Genes with non-standard start/stop codons, internal stop codons, or CDS lengths not divisible by three were excluded. For each species, CDS and protein sequences shorter than 30 residues or longer than 4,000 residues were filtered out.

### 1.5.2 Embedding extraction

CodonBERT embeddings were extracted from the final transformer layer using mean pooling across all token positions, yielding a 768-dimensional vector per CDS. Input sequences were tokenized using CodonBERT’s overlapping codon-pattern tokenizer. ESM-2 (650M parameter variant) embeddings were extracted from the final transformer layer using mean pooling, yielding a 1,280-dimensional vector per protein sequence. DNABERT-2 and Nucleotide Transformer v2 (50M) embeddings were extracted analogously from their respective final layers with model-default tokenization. For each species, embeddings were computed once and stored.

### 1.5.3 Composition residualization

For each gene, a 20-dimensional amino acid composition vector was computed from the translated protein sequence. Both CDS-derived (CodonBERT) and protein-derived (ESM-2) embeddings were regressed against this composition vector using ordinary least squares, and the residuals were retained as composition-orthogonal embeddings. This procedure removes the linear contribution of amino acid identity to both embedding spaces, ensuring that any recovered CDS–protein correspondence reflects features beyond bulk amino acid usage.

### 1.5.4 Paired retrieval with ridge regression

For a given source and target species, gene pairs were split into training and test sets. A ridge regression model (α = 1.0) was trained to map composition-residual CodonBERT embeddings to composition-residual ESM-2 embeddings. For each test query, the predicted ESM-2 embedding was compared against all gallery embeddings via cosine similarity, and R@1 was computed as the fraction of queries for which the true target embedding ranked first. Five independent random seeds (20260627–20260701) were used for all experiments to assess stability. Seed-specific train/test splits are reported in Supplementary Tables.

### 1.5.5 Control conditions

Four control conditions were implemented. (1) *Randomized train pairs*: training pairs were randomly permuted before ridge training. (2) *Synonymous rewrite*: each test CDS was replaced with a synthetic version where every codon was substituted with its highest-frequency synonymous alternative from the species-specific codon usage table; ridge training used the unperturbed CDS as usual. (3) *Codon order shuffle*: codon positions within each CDS were randomly permuted, preserving multiset codon identity but destroying positional context. (4) *Protein gallery shuffle*: ESM-2 gallery embeddings were randomly permuted while keeping queries fixed.

### 1.5.6 Gallery-free per-gene alignment cosine

To avoid gallery-size artifacts in cross-species comparisons, we computed the per-gene alignment cosine: for each orthologous query–target pair, the cosine similarity between the Ridge-predicted ESM-2 embedding (from the source CDS) and the true target ESM-2 embedding was calculated. This metric is computed independently for each gene pair and does not depend on gallery composition or size. Comparisons between species were performed using two-sided Mann-Whitney U tests on the per-gene cosine distributions.

### 1.5.7 Shuffled-pair permutation null

For ortholog-only human↔mouse comparisons, we kept the fitted residual Ridge map and target ESM-2 gallery fixed, then randomly deranged target gallery rows 10,000 times for each seed and direction. The null distribution of mean alignment cosine under random pairing was compared with the observed real-pair mean. Permutation p-values were computed as the fraction of null iterations with mean cosine ≥ observed mean cosine.

### 1.5.8 Evolutionary distance analysis

Orthologous gene pairs between human and each target species (rat, mouse, zebrafish, yeast) were identified via Ensembl Compara. For each species pair, the composition-residual Ridge map was trained and per-gene alignment cosines were computed for all test orthologs. Distributions were compared across species using two-sided Mann-Whitney U tests on the per-gene cosine values, with adjacent species pairs tested sequentially (rat vs. mouse, mouse vs. zebrafish, zebrafish vs. yeast).

### 1.5.9 Model ablation

Embeddings from DNABERT-2 and NT-v2-50M were extracted and composition-residualized identically to the CodonBERT protocol. Ortholog-only human↔mouse per-gene alignment cosines were computed under the same five-seed protocol. Per-gene cosines were averaged across five seeds before summarizing distributions and performing Mann-Whitney U tests.

### 1.5.10 mRNA stability, codon optimality, and functional gradient analyses

mRNA half-life data for human genes were obtained from Tani et al. (2012). CAI was computed using the species-specific codon usage table from the Codon Usage Database. Codon entropy was computed as the Shannon entropy of codon usage frequencies within each CDS. Essential gene annotations were retrieved from the OGEE database; housekeeping and tissue-specific gene annotations were obtained from the Human Protein Atlas. Spearman correlations and Mann-Whitney U tests were computed as noted, with five-seed multiseed analysis used where stated.

### 1.5.11 Data and code availability

All embedding datasets, experiment configurations, and analysis code are available at [repository URL]. Source data for all figures are provided as Supplementary Data files.

## 1.7 Figure Legends

**Supplementary Figure 1.**
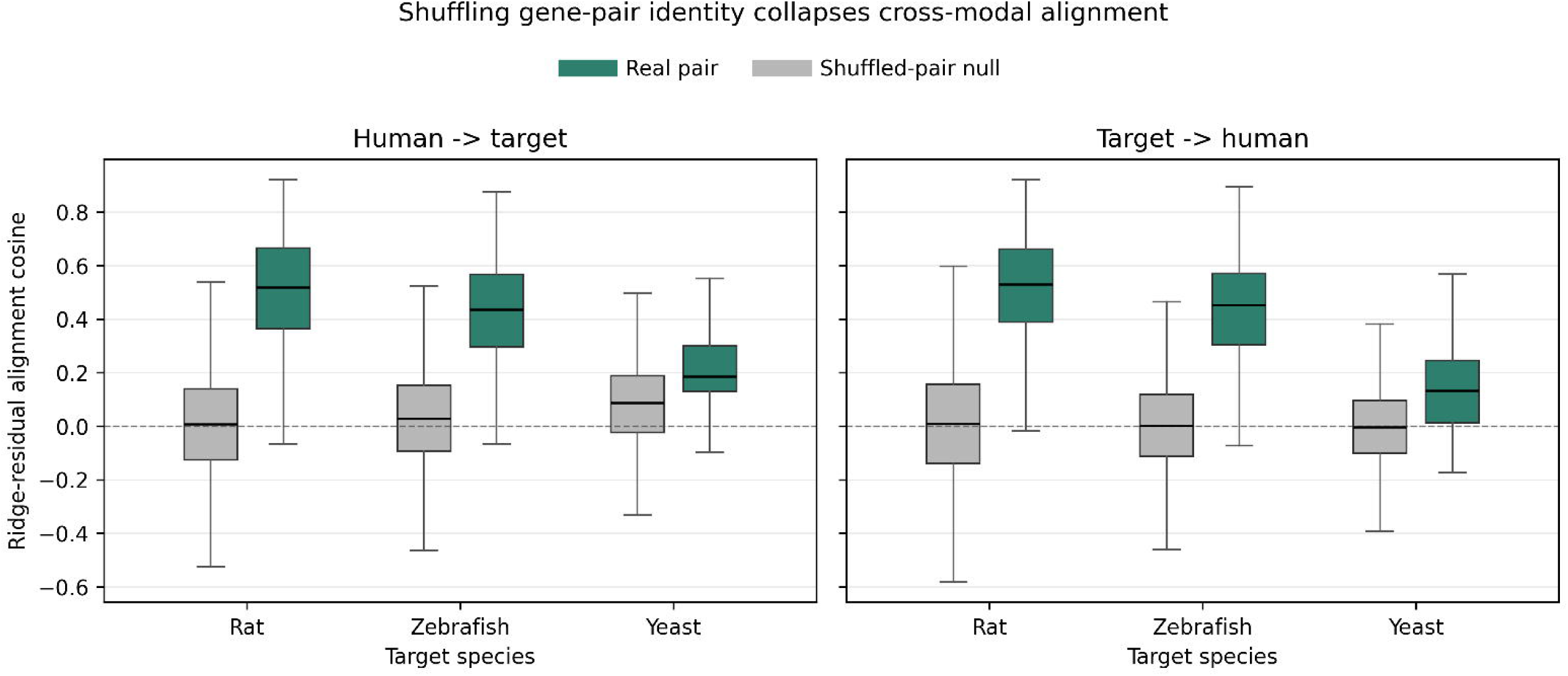
Codon entropy, mRNA stability, and functional gradient analyses. **(a)** Spearman correlations between alignment cosine and mRNA half-life (ortholog-only, 5-seed multiseed). Mean ρ ≈ −0.048, 95% CI [−0.173, 0.076] crosses zero. **(b)** Codon entropy vs. alignment cosine across five seeds. Mean Spearman ρ with 95% CI crossing zero. **(c)** Per-gene hit rates across functional gene categories (essential, housekeeping, tissue-specific). No significant differences detected (Mann-Whitney p > 0.2 for all comparisons).

**Supplementary Figure 2.**
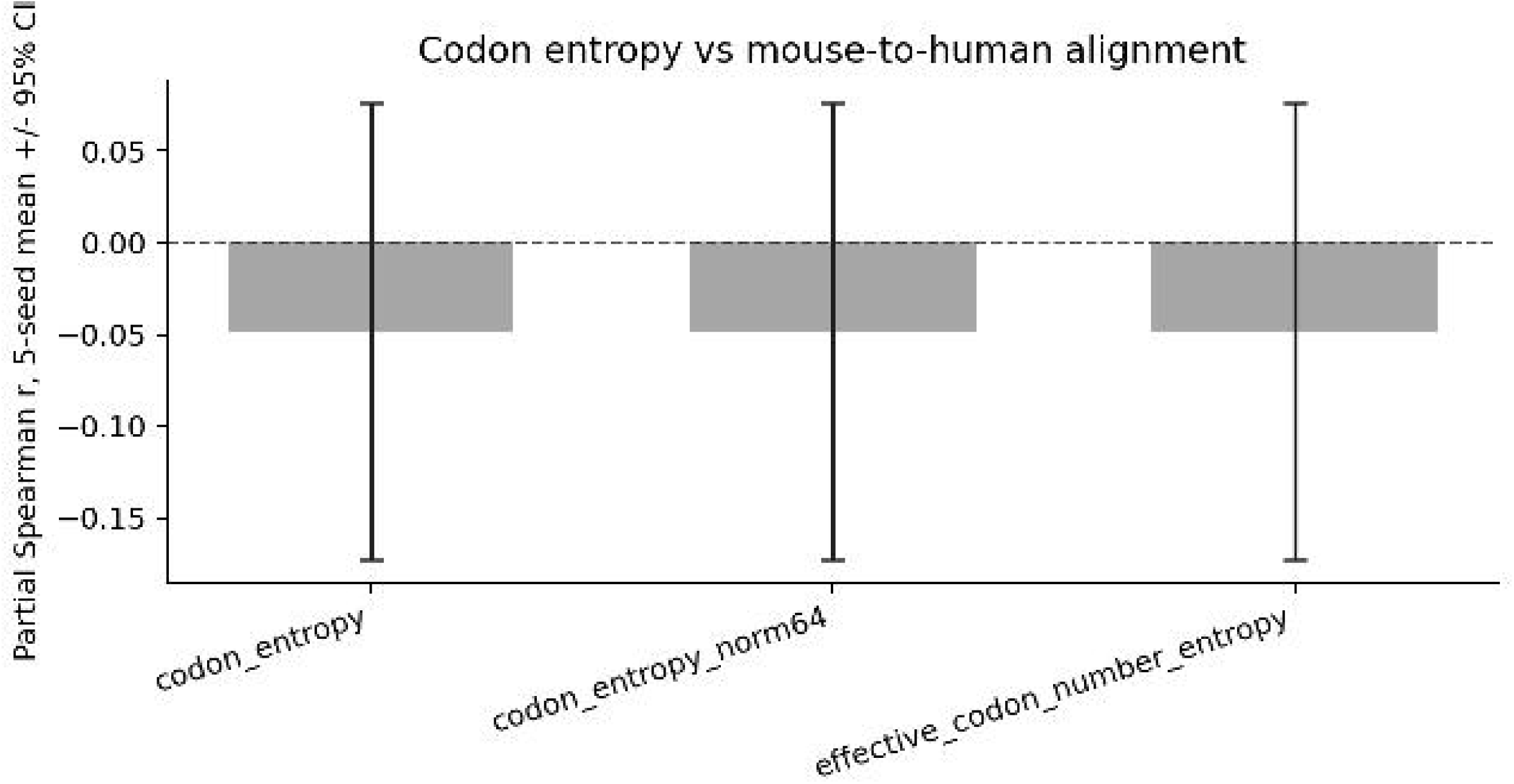
Gallery-size robustness controls for evolutionary distance analysis. R@1 after gallery-size normalization: cap_65 (all galleries capped at 65 genes by random subsampling) and exact_50 (all galleries subsampled to exactly 50 genes). Both procedures confirm rat > mouse > zebrafish ordering. Ten randomization iterations per seed; mean ± SD shown.

## 1.6 Supplementary Tables

See `supplementary_tables_v2.md` for: - Supplementary Table 1: Full within-species and cross-species transfer per-seed results - Supplementary Table 2: Ortholog-only multiseed per-seed detail - Supplementary Table 3: Evolutionary distance per-gene cosine distributions and Mann-Whitney tests - Supplementary Table 4: Shuffled-pair permutation null per-seed results - Supplementary Table 5: Model ablation per-gene cosine and R@1 per-seed results - Supplementary Table 6: mRNA stability, codon optimality, and functional gradient correlations

---

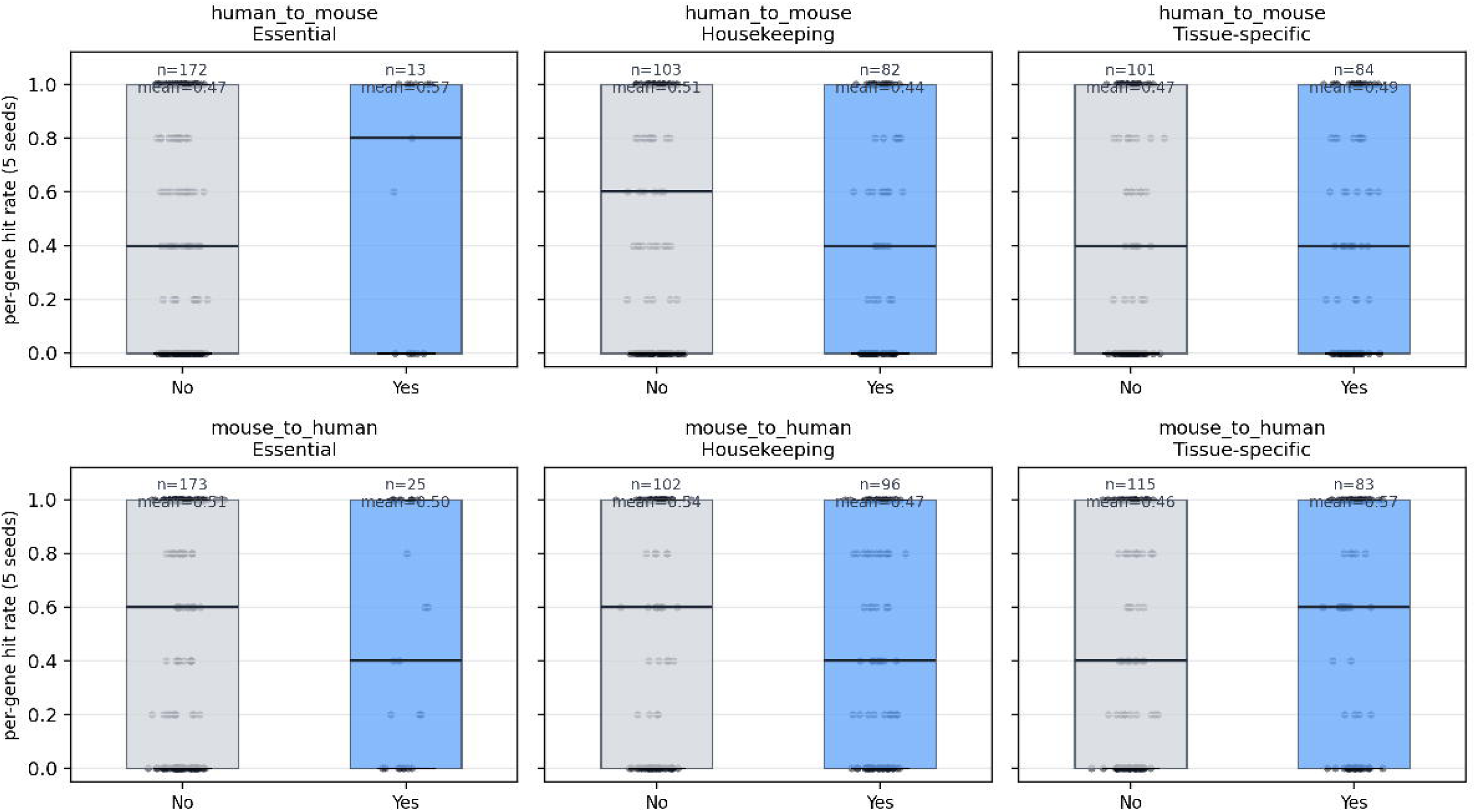

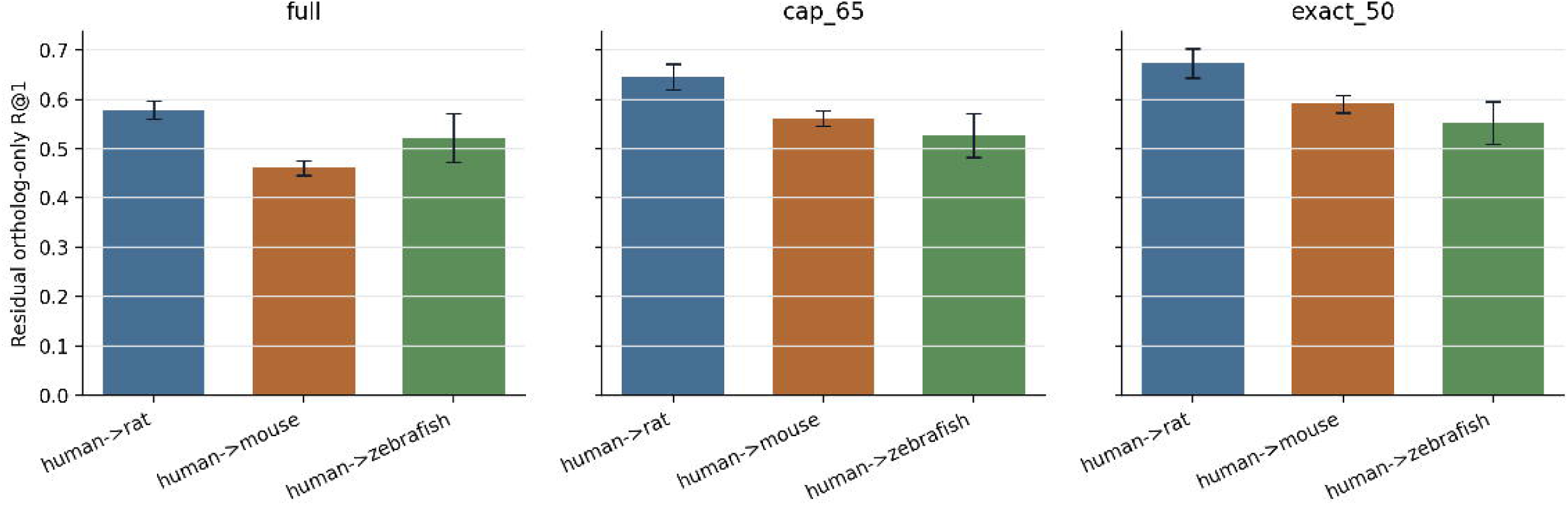

