## Supplementary Tables for "1 CodonBERT and ESM-2 Embedding Spaces Share an Evolutionarily Conserved Paired Geometry Encoding Synonymous Codon Information"

### Supplementary Tables — CodonBERT-ESM Paired Geometry

---

#### Supplementary Table 1: Within-Species and Cross-Species Transfer Per-Seed Results

Human and mouse within-species and cross-species composition-residual R@1 for real pairs and all control conditions, across five seeds (20260627–20260701).

##### Human Within-Species

| Seed | Real | Synonymous Rewrite | Codon Shuffle | Protein Shuffle | Random |
|---|---|---|---|---|---|
| 20260627 | 0.258 | 0.186 | 0.054 | 0.046 | 0.002 |
| 20260628 | 0.275 | 0.191 | 0.059 | 0.050 | 0.000 |
| 20260629 | 0.259 | 0.188 | 0.068 | 0.051 | 0.000 |
| 20260630 | 0.282 | 0.199 | 0.051 | 0.051 | 0.000 |
| 20260701 | 0.258 | 0.172 | 0.058 | 0.049 | 0.001 |
| **Mean** | **0.266** | **0.187** | **0.058** | **0.049** | **0.001** |

##### Mouse Within-Species

| Seed | Real | Synonymous Rewrite | Codon Shuffle | Protein Shuffle | Random |
|---|---|---|---|---|---|
| 20260627 | 0.269 | 0.190 | 0.053 | 0.050 | 0.001 |
| 20260628 | 0.270 | 0.185 | 0.061 | 0.048 | 0.001 |
| 20260629 | 0.262 | 0.186 | 0.058 | 0.048 | 0.000 |
| 20260630 | 0.266 | 0.177 | 0.055 | 0.052 | 0.001 |
| 20260701 | 0.261 | 0.175 | 0.059 | 0.046 | 0.001 |
| **Mean** | **0.266** | **0.183** | **0.057** | **0.049** | **0.001** |

##### Cross-Species Transfer (Full Panel)

| Direction | Real | Synonymous Rewrite | Codon Shuffle | Protein Shuffle | Random |
|---|---|---|---|---|---|
| Human → Mouse | 0.254 | 0.164 | 0.062 | 0.041 | 0.002 |
| Mouse → Human | 0.262 | 0.174 | 0.058 | 0.043 | 0.003 |

---

#### Supplementary Table 2: Ortholog-Only Multiseed Per-Seed Detail

Composition-residual R@1 for ortholog-only human ↔ mouse evaluation.

##### Human → Mouse

| Seed | n_test | Real | Syn Rewrite | Codon Shuffle | Protein Shuffle | Random |
|---|---|---|---|---|---|---|
| 20260627 | 185 | 0.481 | 0.319 | 0.114 | 0.108 | 0.016 |
| 20260628 | 171 | 0.468 | 0.357 | 0.076 | 0.088 | 0.012 |
| 20260629 | 206 | 0.447 | 0.330 | 0.083 | 0.102 | 0.005 |
| 20260630 | 190 | 0.468 | 0.353 | 0.084 | 0.100 | 0.016 |

| Seed | n_test | Real | Syn Rewrite | Codon Shuffle | Protein Shuffle | Random |
|---|---|---|---|---|---|---|
| 20260701 | 193 | 0.440 | 0.337 | 0.109 | 0.150 | 0.010 |
| **Mean** | **189** | **0.461** | **0.339** | **0.093** | **0.110** | **0.012** |

**Mouse → Human**

| Seed | n_test | Real | Syn Rewrite | Codon Shuffle | Protein Shuffle | Random |
|---|---|---|---|---|---|---|
| 20260627 | 198 | 0.510 | 0.389 | 0.146 | 0.101 | 0.010 |
| 20260628 | 192 | 0.505 | 0.349 | 0.193 | 0.120 | 0.000 |
| 20260629 | 186 | 0.575 | 0.398 | 0.194 | 0.108 | 0.011 |
| 20260630 | 174 | 0.546 | 0.402 | 0.138 | 0.115 | 0.000 |
| 20260701 | 181 | 0.552 | 0.409 | 0.160 | 0.094 | 0.006 |
| **Mean** | **186** | **0.538** | **0.389** | **0.166** | **0.107** | **0.005** |

---

#### Supplementary Table 3: Evolutionary Distance Per-Gene Alignment Cosine

Gallery-free per-gene alignment cosine (Ridge-predicted vs. true ESM-2) across species.

**Forward Direction (Human CDS query → target protein)**

| Target | n_genes | Mya | Median | Q10 | Q25 | Q75 | Q90 | Adjacent MW p |
|---|---|---|---|---|---|---|---|---|
| Rat | 631 | 20 | 0.519 | 0.24 | 0.35 | 0.66 | 0.79 | — |
| Mouse | 945 | 90 | 0.466 | 0.21 | 0.33 | 0.61 | 0.72 | $7 \times 10^{7}$ |
| Zebrafish | 321 | 435 | 0.435 | 0.16 | 0.29 | 0.56 | 0.72 | $8 \times 10^{3}$ |
| Yeast | 140 | 1,300 | 0.186 | −0.05 | 0.06 | 0.32 | 0.44 | $4 \times 10^{26}$ |

**Reverse Direction (Target CDS query → Human protein)**

| Source | Median | Q10 | Q25 | Q75 | Q90 | Adjacent MW p |
|---|---|---|---|---|---|---|
| Rat → Human | 0.530 | 0.26 | 0.38 | 0.68 | 0.80 | — |
| Mouse → Human | 0.533 | 0.25 | 0.37 | 0.67 | 0.79 | 0.215 (ns) |
| Zebrafish → Human | 0.452 | 0.18 | 0.30 | 0.59 | 0.73 | $4 \times 10^{9}$ |
| Yeast → Human | 0.210 | −0.04 | 0.07 | 0.34 | 0.46 | $5 \times 10^{41}$ |

---

#### Supplementary Table 4: Shuffled-Pair Permutation Null Per-Seed Results

10,000 random derangements of target gallery rows per seed/direction. Ridge map and gallery distribution held fixed.

| Direction | Seed | n | Real Mean | Real Median | Null Mean | Null 95% CI | Δ Mean | Permutation p |
|---|---|---|---|---|---|---|---|---|
| Human → Mouse | 20260627 | 185 | 0.470 | 0.467 | 0.008 | [−0.019, 0.036] | 0.462 | $<10^{4}$ |
| Human → Mouse | 20260628 | 171 | 0.445 | 0.444 | 0.007 | [−0.019, 0.034] | 0.438 | $<10^{4}$ |

| Direction | Seed | n | Real Mean | Real Median | Null Mean | Null 95% CI | Δ Mean | Permutation p |
|---|---|---|---|---|---|---|---|---|
| Human→Mouse | 20260629 | 206 | 0.452 | 0.459 | 0.013 | [−0.013, 0.037] | 0.439 | $<10^{4}$ |
| Human→Mouse | 20260630 | 190 | 0.474 | 0.479 | 0.003 | [−0.024, 0.030] | 0.470 | $<10^{4}$ |
| Human→Mouse | 20260701 | 193 | 0.458 | 0.468 | 0.009 | [−0.018, 0.036] | 0.448 | $<10^{4}$ |
| Mouse→Human | 20260627 | 198 | 0.498 | 0.513 | 0.007 | [−0.020, 0.034] | 0.491 | $<10^{4}$ |
| Mouse→Human | 20260628 | 192 | 0.509 | 0.536 | 0.011 | [−0.017, 0.039] | 0.499 | $<10^{4}$ |
| Mouse→Human | 20260629 | 186 | 0.529 | 0.545 | 0.003 | [−0.025, 0.032] | 0.525 | $<10^{4}$ |
| Mouse→Human | 20260630 | 174 | 0.499 | 0.526 | 0.005 | [−0.026, 0.035] | 0.494 | $<10^{4}$ |
| Mouse→Human | 20260701 | 181 | 0.496 | 0.542 | 0.002 | [−0.026, 0.030] | 0.495 | $<10^{4}$ |
| **H→M Mean** | | **189** | **0.460** | **0.463** | **0.008** | | **0.452** | **$<10^{4}$** |
| **M→H Mean** | | **186** | **0.506** | **0.532** | **0.005** | | **0.501** | **$<10^{4}$** |

---

#### Supplementary Table 5: Model Ablation — Per-Gene Alignment Cosine

Per-gene alignment cosine (five-seed averaged) for ortholog-only human↔mouse comparison across three nucleotide language models.

**Human→Mouse**

| Model | n_genes | Median | Q25 | Q75 | Mann-Whitney vs CodonBERT |
|---|---|---|---|---|---|
| CodonBERT | 626 | 0.467 | 0.327 | 0.613 | — |
| DNABERT-2 | 626 | 0.275 | 0.145 | 0.404 | $p = 2 \times 10^{49}$ |
| NT-v2-50M | 626 | 0.259 | 0.113 | 0.392 | $p = 2 \times 10^{58}$ |

**Mouse→Human**

| Model | n_genes | Median | Q25 | Q75 | Mann-Whitney vs CodonBERT |
|---|---|---|---|---|---|
| CodonBERT | 638 | 0.526 | 0.379 | 0.654 | — |
| DNABERT-2 | 638 | 0.309 | 0.161 | 0.447 | $p = 1 \times 10^{55}$ |
| NT-v2-50M | 638 | 0.281 | 0.152 | 0.398 | $p = 7 \times 10^{77}$ |

**Ortholog-Only R@1 Comparison**

| Model | Human→Mouse R@1 | Mouse→Human R@1 |
|---|---|---|
| CodonBERT | 0.461 | 0.538 |
| DNABERT-2 | 0.161 | — |
| NT-v2-50M | 0.129 | — |

| Model | Human→Mouse R@1 | Mouse→Human R@1 |
|---|---|---|

#### Supplementary Table 6: mRNA Stability, Codon Optimality, and Functional Gradient Correlations

| Analysis | Metric | Result | 95% CI | Significance |
|---|---|---|---|---|
| Codon entropy (5-seed) | Spearman ρ | −0.048 | [−0.173, 0.076] | ns |
| mRNA half-life (5-seed) | Spearman ρ | — | [crossing zero] | ns (4/5 seeds) |
| CAI | Spearman ρ | ~0.01 | — | $p = 0.76$ |
| GC3 | Spearman ρ | ~0.02 | — | $p = 0.61$ |
| CDS length | Spearman ρ | ~0.13 | — | $p < 0.001$ |
| Essential vs non-essential | Mann-Whitney | — | — | $p > 0.2$ |
| Housekeeping vs non-housekeeping | Mann-Whitney | — | — | $p > 0.2$ |
| Tissue-specific vs broad | Mann-Whitney | — | — | $p > 0.2$ |
